## Supplementary Figures for "CRISPR screen decodes SWI/SNF chromatin remodeling complex assembly"

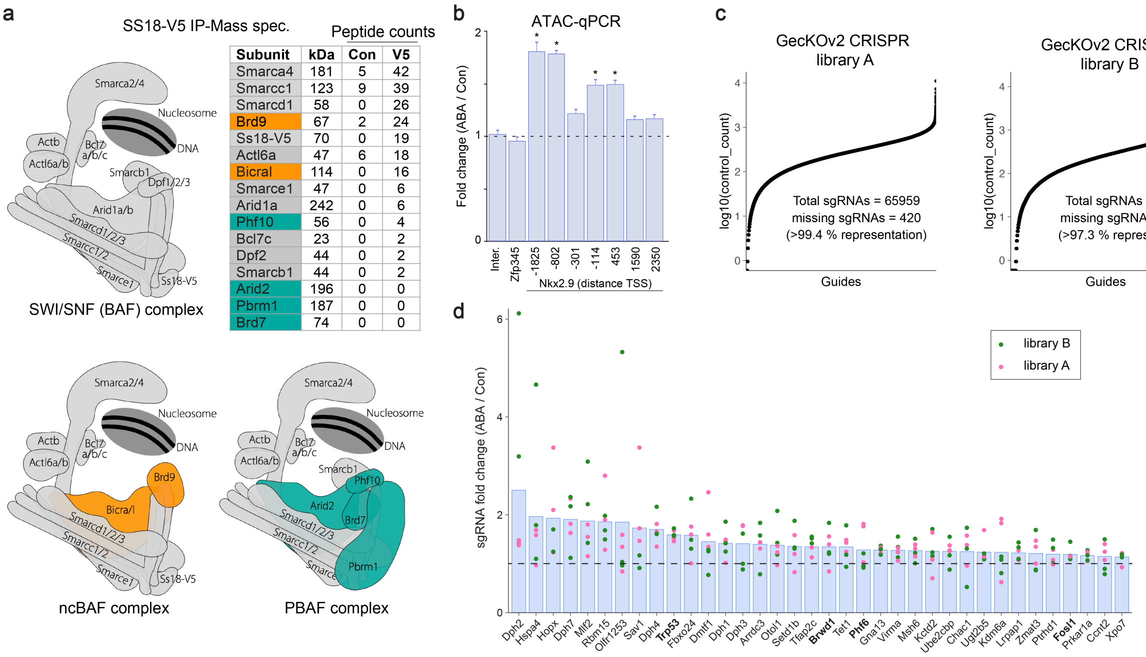


| Subunit | kDa | Con | V5 |
|---|---|---|---|
| Smarca4 | 181 | 5 | 42 |
| Smarcc1 | 123 | 9 | 39 |
| Smarcd1 | 58 | 0 | 26 |
| Brd9 | 67 | 2 | 24 |
| Ss18-V5 | 70 | 0 | 19 |
| Actl6a | 47 | 6 | 18 |
| Bicral | 114 | 0 | 16 |
| Smarce1 | 47 | 0 | 6 |
| Arid1a | 242 | 0 | 6 |
| Phf10 | 56 | 0 | 4 |
| Bcl7c | 23 | 0 | 2 |
| Dpf2 | 44 | 0 | 2 |
| Smarcb1 | 44 | 0 | 2 |
| Arid2 | 196 | 0 | 0 |
| Pbrm1 | 187 | 0 | 0 |
| Brd7 | 74 | 0 | 0 |

### Supplementary Fig 1 | Related to Figure 1

**a**, Peptide counts from V5 IP-MS in WT and V5-SS18 expressing mESCs. Shown are the subunits from the different SWI/SNF complexes expressed in mESCs. As expected only SWI/SNF (grey) and ncBAF (orange) are pulled-down via the SS18 subunit, whereas PBAF (green) subunits are not enriched in the V5-IP.
**b**, ATAC-qPCR shows a gain in chromatin accessibility at *Nkx2.9* locus after ABA induced SWI/SNF-recruitment for 24 hours. qPCR analysis performed using seven primer pairs tilling the *Nkx2.9* locus and three control primers (*Intergenic*, *Zfp345* and *Rpl12*). Fold change normalized to *Rpl12*. Statistical significance was calculated using t-test (* = $p < 0.05$), replicates n = 4.
**c**, Distribution of sgRNAs in library A and B in control cells after GecKOv2 lentivirus library transduction. Sequencing reveals diverse representation and high coverage of sgRNAs.
**d**, Fold change of sgRNAs in library A and B. Bars represent the mean fold change between ABA treated and EtOH controls. Depicted genes have MAGeCK Enrichment score < 1.00E-4, except for Fosl1 (E. score = 1.53E-4), Kdm6a (E. score = 5.99E-3) and Setd1b (E. score = 2.96E-4). Genes in bold are previously identified interactors of the SWI/SNF complex. n = 6 sgRNAs targeting all mouse genes.

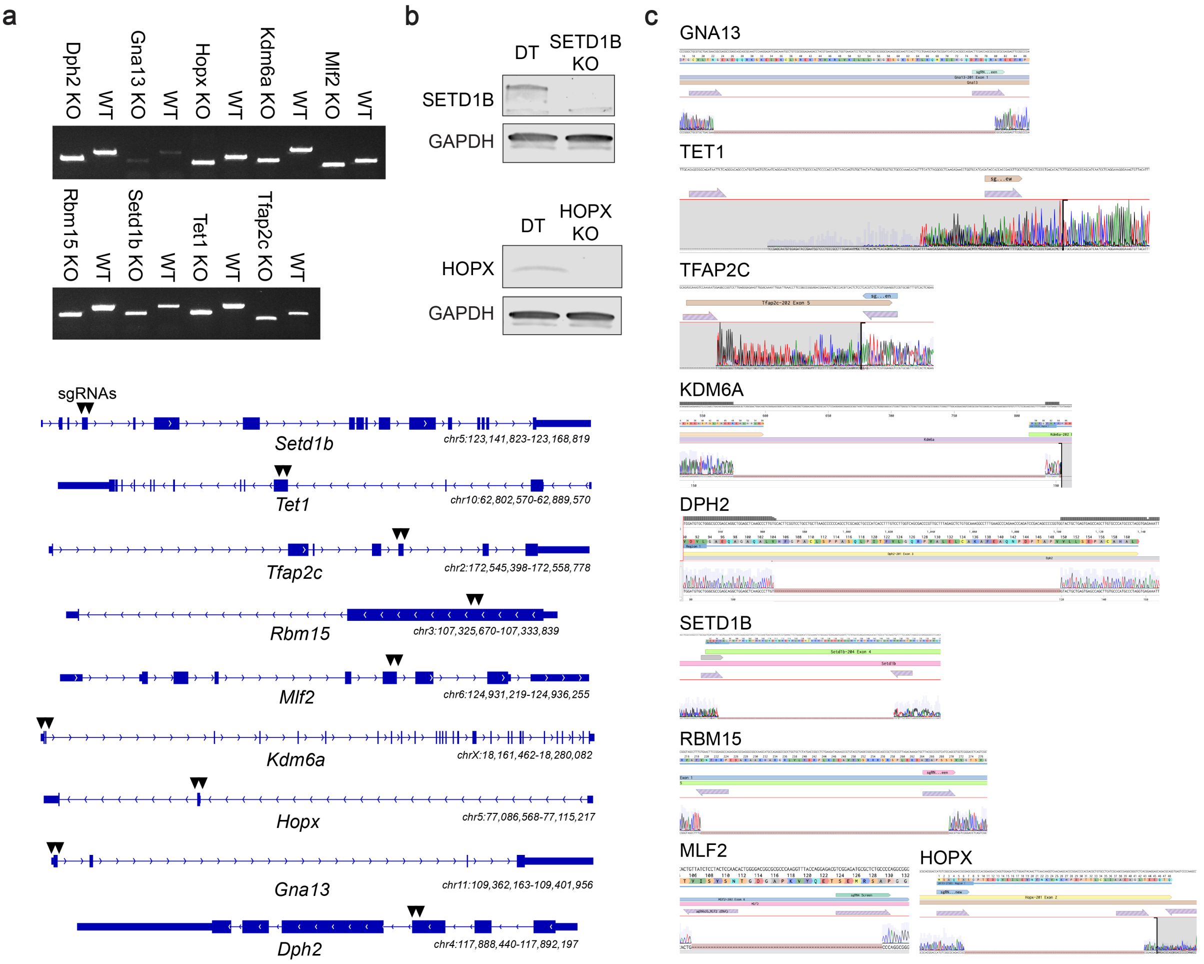


### Supplementary Fig 2 | Related to Figure 2

**a**, Genotyping PCRs of CRISPR KO mESC lines compared to WT mESCs. Shifted bands indicate successful KO of the target locus. Below, schematic of *Dph2*, *Mlf2*, *Rbm15*, *Hopx*, *Tfap2c*, *Gna13*, *Setd1b*, *Kdm6a* and *Tet1* genomic loci in mouse genome (mm10); black arrows indicate exons targeted by pairs of gRNAs to KO individual genes.
**b**, Western blot analysis of SETD1B and HOPX KO mESC clones. Antibodies ordered for TET1, KDM6A and DPH2 did not work. GAPDH as loading control.
**c**, Sanger sequencing analysis shows KO by indel/frameshift of all 9 genes in individual mESC clones.

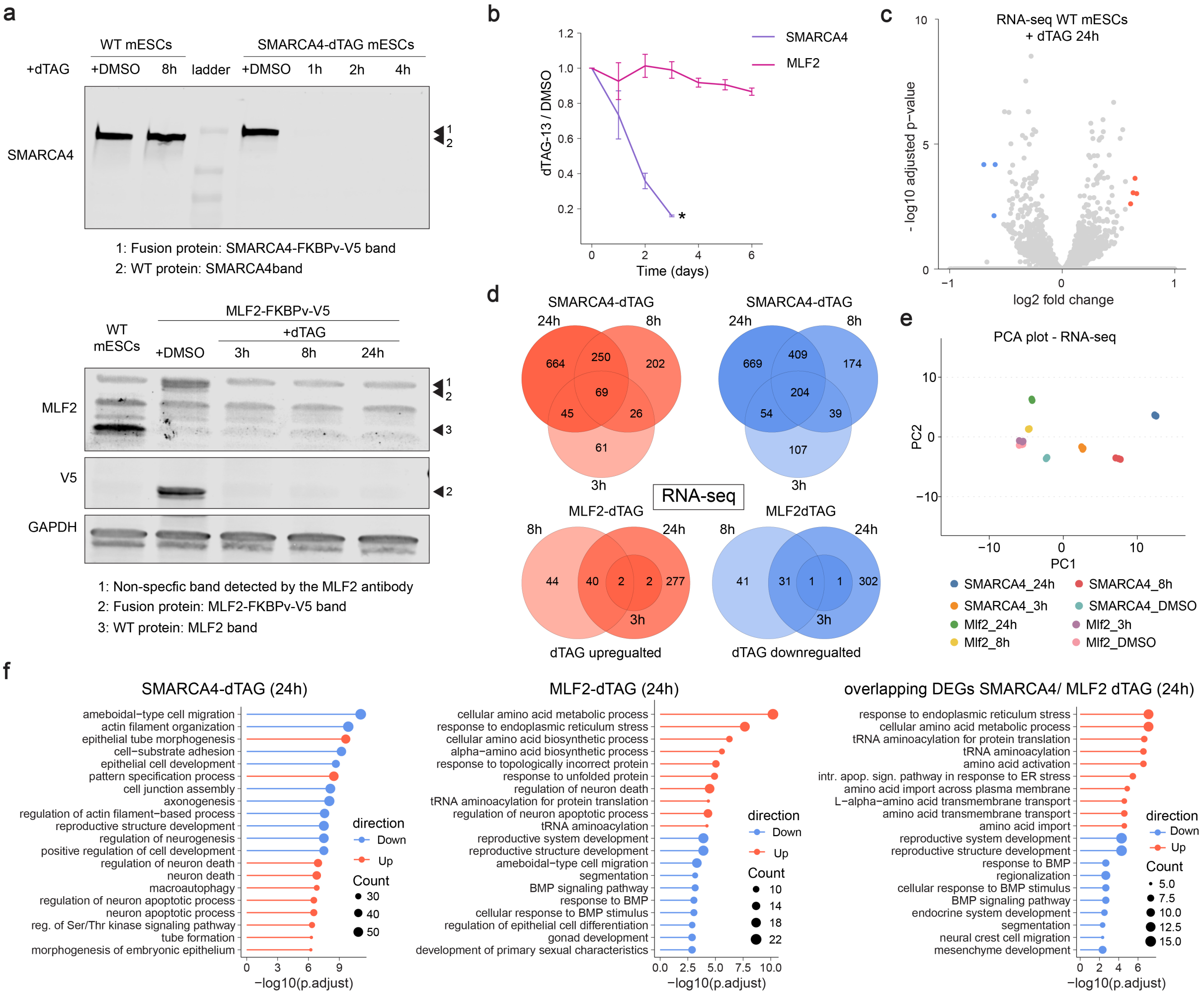


### Supplementary Fig 3 | Related to Figure 3

**a**, Top: Western blot analysis of SMARCA4 levels in SMARCA4-dTAG mESCs treated with DMSO, 1h, 2h and 4h with dTAG-13 and WT mESCs treated with DMSO and dTAG-13 for 8h. GAPDH as loading control. Bottom: Western blot analysis of MLF2 and V5 levels in MLF2-dTAG mESCs treated with DMSO, 3h, 8h and 24h with dTAG-13 compare to WT mESCs. GAPDH as loading control. Note: MLF2 antibody detects a non-specific band just above the MLF2-FKBPv-V5 band.
**b**, Growth curve of MLF2-dTAG and SMARCA4-dTAG mESCs treated with dTAG-13. Replicates n = 3, * indicates complete cell death.
**c**, Volcano plot shows only 7 DEGs in WT mESC after 24h of dTAG-13 treatment compared to DMSO controls. Significantly downregulated genes are colored in blue and upregulated in red (sig. = padj. $< 0.05$ & $|FC| > 1.5$; replicates n = 3).
**d**, Venn diagrams showing overlaps of DEGs from RNA-seq at 3h, 8h and 24h of dTAG-13 treatment in MLF2 and SMARCA4 dTAG treated mESCs. Upregualted genes are shown in red and downregulated genes are shown in blue.
**e**, PCA plot of RNA-seq samples from MLF2-dTAG and SMARCA4-dTAG mESCs treated with dTAG13 (3h, 8h, 24h) or DMSO. Replicates n = 3.
**f**, GO-terms of DEGs from SMARCA4 dTAG and MLF2 dTAG cells after 24h of dTAG-13 treatment.

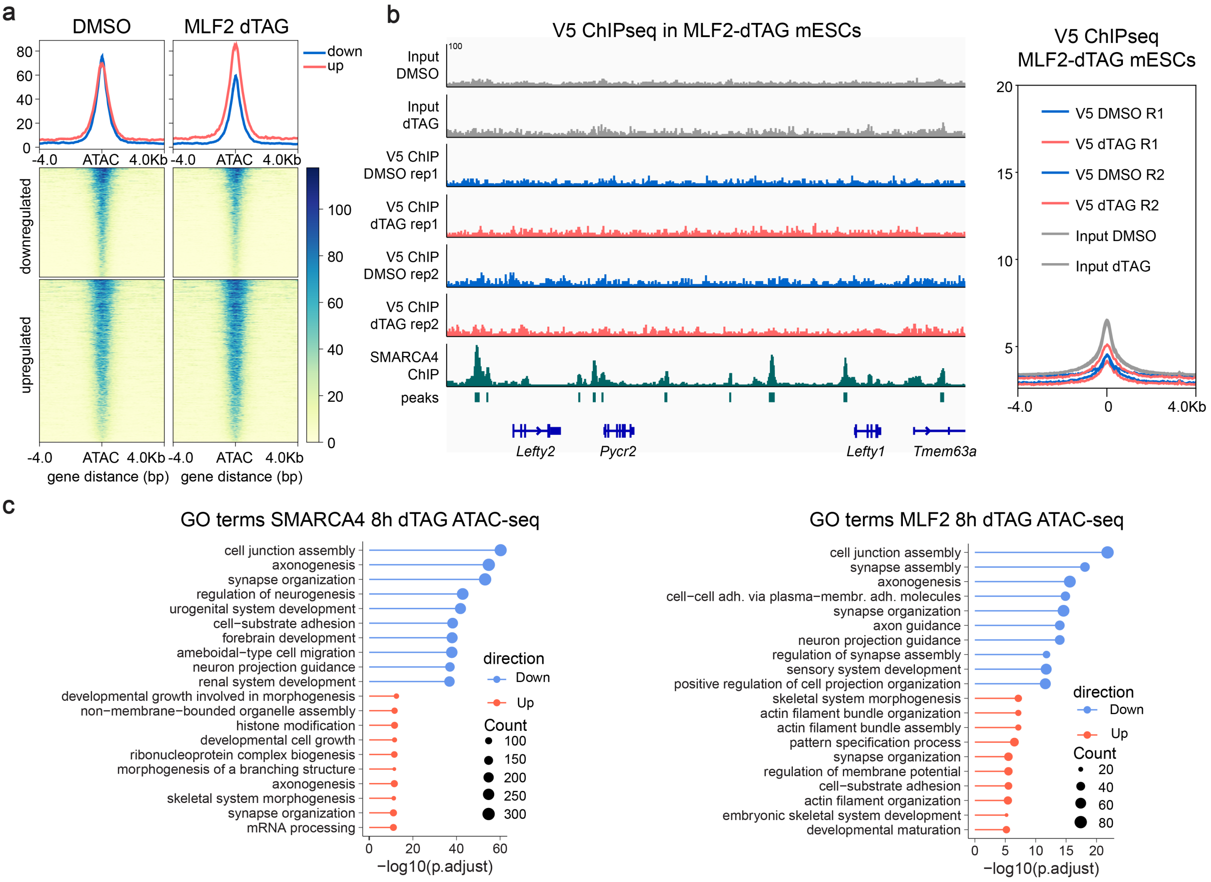


### Supplementary Fig 4 | Related to Figure 4

**a**, ATAC-seq signal at upregulated and downregulated peaks in MLF2-dTAG mESCs treated with dTAG-13 for 8h compared to DMSO control condition.
**b**, Genome browser tracks depicting V5 ChIP-seq and input signal in MLF2-dTAG mESCs compared to published SMARCA4 ChIP-seq[47] peaks at the *Lefty1* locus (left). V5 ChIP-seq signal compared to input signal shows no enrichment and suggests MLF2 does not bind chromatin (right).
**c**, GO-terms of genes associated to changed ATAC-seq peaks in SMARCA4-dTAG and MLF2-dTAG mESCs after 8h of dTAG-13 treatment.

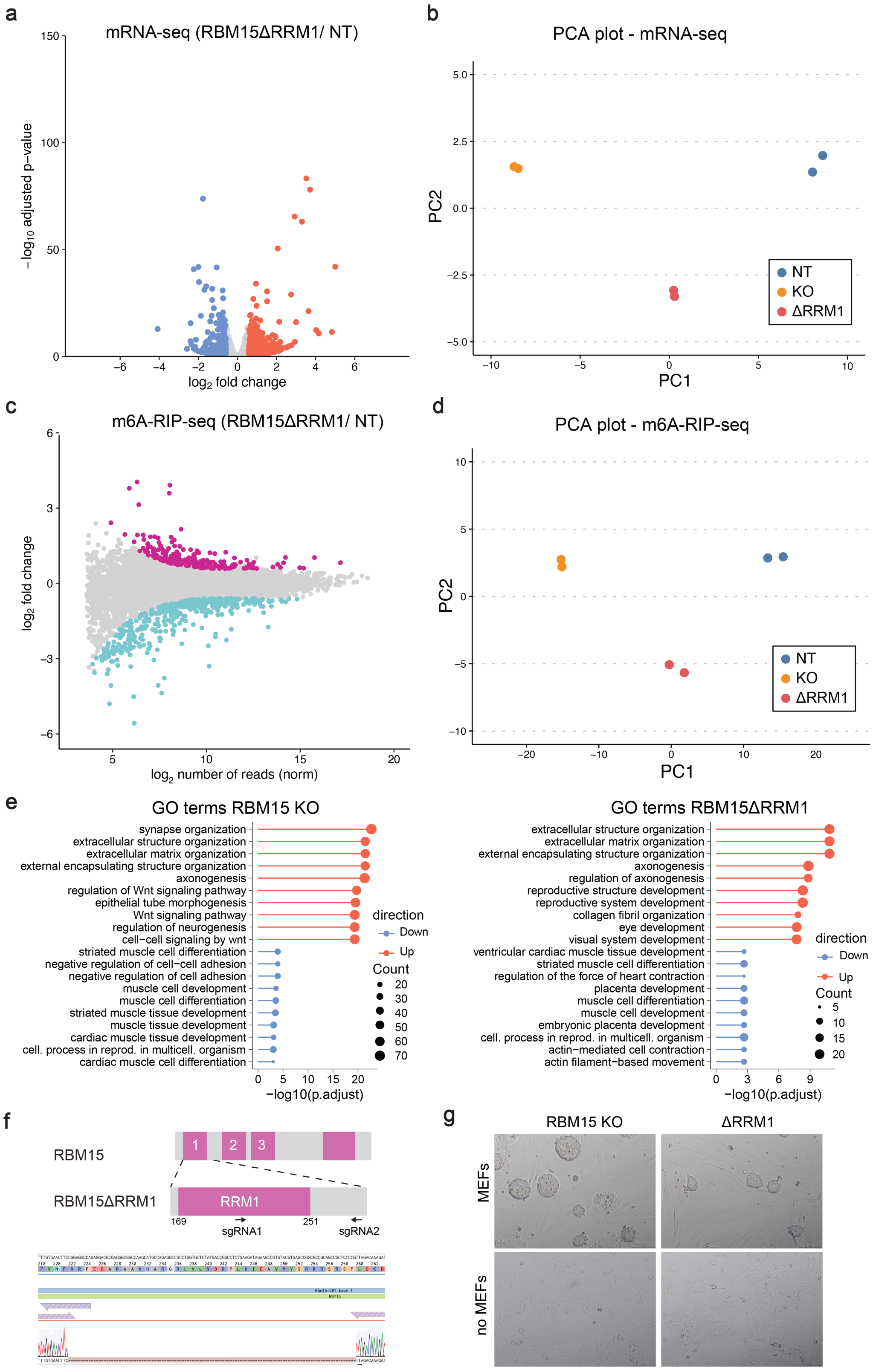


### Supplementary Fig 5 | Related to Figure 5

**a,** Volcano plot showing differentially expressed genes (DEGs) between NT and RBM15 ΔRRM1 mutant mESCs (n=2, FDR < 0.05 and |FC| > 1.5). The 265 downregulated genes are labeled in blue, the 253 upregulated genes are in red.
**b**, Plot shows principal component analysis (PCA) for RNA-seq from NT, RBM15 KO and ΔRRM1 mESCs.
**c**, MA plot showing differentially m6A modified mRNA peaks between NT and RBM15 ΔRRM1 mutant mESCs (n=2, FDR < 0.05 and |FC| > 1.5). The 232 downregulated peaks are labeled in cyan, the 423 upregulated peaks are in magenta.

**d,** Plot shows principal component analysis (PCA) for m6A-RIP-seq from NT, RBM15 KO and ΔRRM1 mESCs.
**e**, GO-term analysis of up- and down-regulated genes in RBM15 KO and ΔRRM1 mESCs.
**f,** Schematic of *Rbm15* ΔRRM1 disruption strategy in mESCs; black arrows indicate pairs of sgRNAs used to target RRM1. Sanger sequencing analysis confirms in-frame deletion and disruption of RRM1 domain of RBM15.
**g,** Images of RBM15 KO and RBM15 ΔRRM1 mESCs grown on MEF-feeders and after extended culture in feeder-free conditions (no MEFs).

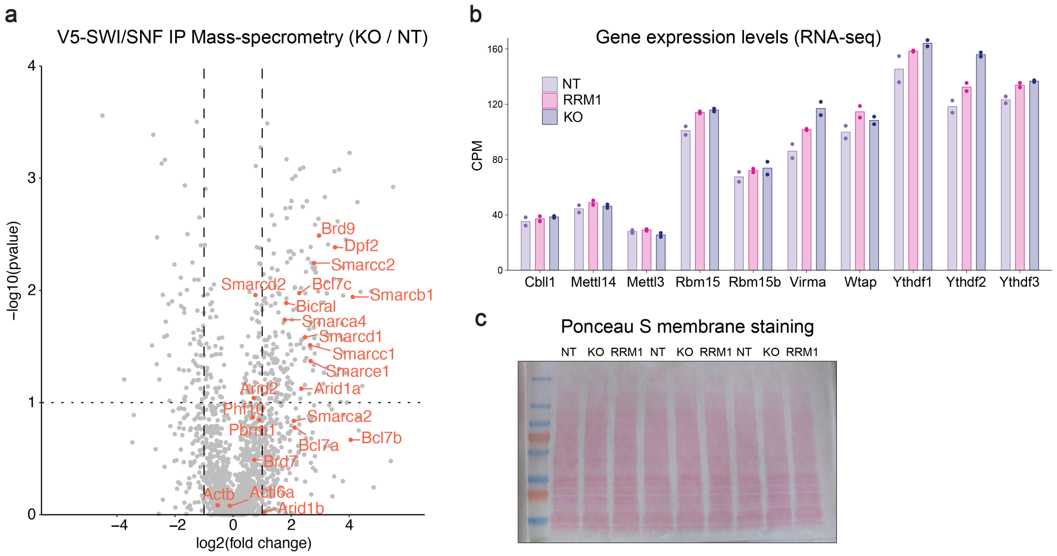


### Supplementary Fig 6 | Related to Figure 6

**a**, Volcano plot showing all proteins detected in the V5-SWI/SNF IP-MS proteomics analysis. Fold change reflects protein enriched in RBM15 KO mESCs compared to NT mESCs. Highlighted in red are the SWI/SNF subunit proteins.
**b**, RNA-seq counts for genes in the m6A RNA methylation pathway in NT, RBM15 KO and ΔRRM1 mESCs.
**c**, Ponceau-S staining of western blot membrane shows equal loading of protein lysates for analysis of SWI/SNF subunit levels in NT, RBM15 KO and ΔRRM1 mESCs.

**Supplementary tables:**

**Supplementary table 1.** MAGeCK analysis of enrichment scores for all mouse genes in CRISPR KO screen of *DT-ZF-Nkx2.9* mESCs

**Supplementary table 2.** DESeq2 analysis of differentially expressed genes from RNA-seq experiments in MLF2-dTAG and SMARCA4-dTAG mESCs treated with dTAG13 for 3h, 8h and 24h.

**Supplementary table 3.** DESeq2 analysis of differentially accessible peaks from ATAC-seq experiments in MLF2-dTAG and SMARCA4-dTAG mESCs treated with dTAG13 for 8h.

**Supplementary table 4.** DESeq2 analysis of differentially expressed genes and differential $m^6A$ peaks from RNA-seq and $m^6A$-RIP-seq experiments in NT, RBM15 KO and ΔRRM1 mESCs.

**Supplementary table 5.** Peptide counts and statistical analysis from SWI/SNF IP-MS analysis of NT and RBM15 KO mESCs

**Supplementary table 6**. List of primers and antibodies used in this study.
